# Multi-Model Machine Learning Consensus Identifies a Dual-Interferon Gene Signature in Ischemic Stroke: An Integrative Single-Cell Transcriptomic Analysis

**DOI:** 10.64898/2026.09.25.754010

**Authors:** Alisha Shamim, Angana Chakroborty

## Abstract

**Background:** An Ischemic stroke is one of the main causes of death and long-term disability around the world. There are very few treatment options available, especially for the cases where the short time period to give clot dissolving medicine has passed (Powers et al., 2019). While single-cell RNA sequencing (scRNA-seq) has emerged as a powerful tool for understanding cell-to-cell differences within a tissue (cellular heterogeneity) (Tang et al., 2009). However translating these high-dimensional datasets into clinically useful biomarkers remains challenging (Rajkomar et al., 2019). A major problem with current biomarker studies is their reliance on a single machine learning model, where results may reflect model bias rather than true biological signals (Lopez-Rincon et al., 2020).

**Methods:** We developed a multi-model consensus framework integrating eight different machine learning methods from three groups: traditional models (logistic regression, random forest, XGBoost), a hybrid deep learning model (CNN-XGBoost), and transformer-inspired models (scGPT, scBERT, sc-Former, Geneformer). The proposed framework was applied to scRNA-seq data from a mouse middle cerebral artery occlusion (MCAO) model (GSE174574; n=6 mice, 54,599 cells). To prevent data leakage, we implemented rigorous subject-wise train/test splitting, ensuring all cells from individual animals remained either in the training sets or in the test sets.

**Results:** All eight models achieved excellent performance on the independent test data, with AUC scores between 0.9778 and 0.9975. From six models that allowed gene-level importance extraction, we identified 63 consensus genes (threshold: *≥*4/6 models), with 17 genes selected by all 6 models. Pathway enrichment analysis showed significant enrichment of interferon signaling, with a novel “dual-interferon” pattern. Twelve of the consensus genes respond to interferon, and five of them (Ifitm6, Ifitm1, Gbp2, Igtp, Tgtp1) respond to both Type I (IFN-*α*/*β*) and Type II (IFN-*γ*) interferons. Notably the consensus included Il1rn (interleukin-1 receptor antagonist), the target of Anakinra—an FDA-approved drug that has completed Phase II stroke trials with promising safety profiles (Smith et al., 2018; Emsley et al., 2005).

**Conclusions:** The Multi-model consensus framework, proposed in this paper, provides a robust and reliable approach to biomarker discovery that goes beyond the limitations of any single method and reduces individual algorithm bias. The convergence of classical, ensemble, and deep learning methods on interferon and IL-1 pathway genes supports their biological validity as stroke biomarkers. Our findings provide a prioritized gene list for further clinical validation and highlight interferon signaling as a possible treatment strategy warranting further investigation.

## 1 Introduction

Stroke is the second leading cause of death globally and a major contributor to adult disability, affecting approximately 12.2 million people annually and causing 6.5 million deaths worldwide (Virani et al., 2021). Ischemic stroke, caused when an artery supplying blood to the brain gets blocked, accounts for about 87% of all stroke cases. It is also responsible for a complex series of cellular and molecular changes that influence patient recovery and survival (Virani et al., 2021).

Despite decades of research, treatment options remain limited. Intravenous thrombolysis with tissue plasminogen activator (tPA) has historically been the primary FDA-approved pharmacological treatment. However, only 3–5% of patients receive it within the recommended time window due to narrow eligibility criteria and contraindications (Hacke et al., 2008; Powers et al., 2019). Mechanical thrombectomy has since extended reperfusion therapy to patients with large-vessel occlusion, but its benefit is very time-critical (Goyal et al., 2016; Saver et al., 2016). Beyond reperfusion, more than a thousand candidate neuroprotective agents have failed to translate from preclinical models into clinical benefit (O’Collins et al., 2006), and no molecular biomarker is yet in routine use for diagnosis, prognosis, or treatment selection (Jickling and Sharp, 2011). This translational gap motivates the search for robust, mechanistically grounded molecular targets.

The development of single-cell RNA sequencing (scRNA-seq) has revolutionized our understanding of the different cell types (Tang et al., 2009). Unlike bulk transcriptomic approaches, scRNA-seq enabled the study of gene expression at the level of individual cells, allowing to identify cell-type–specific patterns that are hidden in tissue-level analyses. In ischemic stroke, scRNA-seq studies have identified different activation states of microglia (Hammond et al., 2019), entry of blood-derived immune cells into the injured brain (Gelderblom et al., 2009; Iadecola et al., 2020), and reactive astrocyte subtypes that play roles in both harmful (neurotoxic) and protective (neuroprotective) brain responses (Liddelow et al., 2017; Zheng et al., 2022). However translating these rich datasets into clinically useful biomarkers remains a significant challenge.

Machine learning (ML) approaches offer powerful computational tools for extracting meaningful patterns from high-dimensional biological data (Rajkomar et al., 2019). Classical algorithms such as logistic regression and random forests provide interpretable models with well-understood statistical properties (Breiman, 2001; Hastie et al., 2009), while gradient boosting methods like XGBoost offer strong predictive performance by combining multiple models (Chen and Guestrin, 2016).

More recently, transformer architecture originally developed for natural language processing (Vaswani et al., 2017) have been adapted for single-cell genomics, with foundation models such as scGPT (Cui et al., 2024), scBERT (Yang et al., 2022), and Geneformer (Theodoris et al., 2023) demonstrating promising results in identifying cell types, predicting gene perturbations, and disease classification. A critical limitation of current computational biomarker studies is their reliance on single ML models. Different algorithmic families focus on different aspects of the data: linear models capture additive effects and provide coefficient-based interpretation; tree-based models excel at detecting nonlinear interactions and offer built-in measures of feature importance; and deep learning models learn complex hierarchical representations but often act as “black boxes” (LeCun et al., 2015; Rudin, 2019). As a result, genes identified by any single model may reflect algorithmic biases as much as biological truth (Strobl et al., 2007; Nogueira et al., 2018), which limits reliability and clinical usefulness of computational findings.

To address this fundamental limitation, we have developed a multi-model consensus framework that uses a diverse set of methods to identify robust biomarker candidates. We reasoned that genes consistently selected by different models with varying learning patterns are more likely to represent true biological signals rather than model-related artifacts—the same principle that underlies ensemble feature selection for biomarker discovery (Dietterich, 2000; Abeel et al., 2010). Here, we applied the proposed framework to scRNA-seq data from a mouse middle cerebral artery occlusion (MCAO) model, integrating eight ML models spanning classical, hybrid, and transformer-inspired architectures. Our analysis identified 63 consensus genes, including a novel dual-interferon pattern and IL-1 pathway genes, providing a prioritized list for translational stroke research.

## 2 Materials And Methods

### 2.1 Data Acquisition and Study Design

We analyzed publicly available single-cell RNA sequencing data from mouse brain tissue after transient middle cerebral artery occlusion (tMCAO), obtained from the Gene Expression Omnibus database (GEO accession: GSE174574) (Zheng et al., 2022). The dataset comprises brain tissue from six adult C57BL/6 mice: three sham-operated controls and three animals that underwent 60 minutes of transient MCAO by the intraluminal filament method (Longa et al., 1989) followed by 24 hours of reperfusion. Brain tissue was separated into individual cells and processed using the 10x Genomics Chromium platform (Zheng et al., 2017) for single-cell capture and library preparation.

### 2.2 Quality Control and Preprocessing

Raw count matrices were processed using Seurat v5 (Butler et al., 2018; Hao et al., 2021) following standard quality control (QC) protocols for scRNA-seq data (Luecken and Theis, 2019). We applied the following filtering criteria to remove low-quality cells and possible doublets (Ilicic et al., 2016):

- Minimum genes detected per cell: 200
- Maximum genes detected per cell: 2,500 (to remove possible doublets)
- Maximum mitochondrial gene percentage: 10% (to remove dying cells)
- Minimum cells expressing each gene: 3 (to remove rarely expressed genes)

We took these threshold values from the standard Seurat workflow (Luecken and Theis, 2019) and then checked them against our own data. Supplementary Figure S1 shows the distribution of each metric with the cut-off lines marked. The median number of genes per cell was 1,324, with a long tail of cells above this. The 2,500 limit sits above the 95th percentile and removes 2,634 cells (4.5%) from the top of this tail. This is where doublets are found, because two cells captured in one droplet show about twice the normal number of genes. For the mitochondrial threshold we used 10% instead of the 5% used in the standard workflow. Osorio and Cai (2021) showed that the right cut-off depends on the tissue and that 5% does not suit every dataset. In our data, a 5% cut-off would remove 12.6% of all cells, while 10% removes only 1,299 cells (2.2%) at the top of the distribution (Supplementary Figure S1C). Brain cells naturally have a higher mitochondrial fraction because of their high energy use. This is not caused by the stroke, since control and MCAO cells have the same median value (2.4% and 2.3%). Nine cells failed both criteria, so 3,924 cells were removed in total.

Removing genes that are detected in very few cells is a standard step in scRNA-seq quality control (Luecken and Theis, 2019). We used the usual minimum of three cells. This removes any gene detected in fewer than three of the 58,523 cells. A gene found in only one or two cells cannot help in any comparison between groups, and such low counts usually come from ambient RNA or mapping errors rather than real expression. This filter reduced the gene set from 27,998 to about 18,676 without losing any information useful to the models.

We did not apply dedicated doublet-detection tools (McGinnis et al., 2019; Wolock et al., 2019) or ambient-RNA correction (Young and Behjati, 2020) in this study. These might be considered for future enhancements of the pipeline.

After QC filtering, the dataset comprised 54,599 high-quality cells: 26,062 cells from sham-operated controls and 28,537 cells from MCAO-treated mice. Data were normalized using the LogNormalize method (scale factor = 10,000), which applies a log-transformation to the library-size normalized counts (counts normalized for total reads per cell) to stabilize variance (Luecken and Theis, 2019). We identified 3,000 highly variable genes (HVGs) using the variance stabilizing transformation (VST) method (Stuart et al., 2019), which describes the mean-variance relationship to select genes with high biological variability relative to their expression level.

### 2.3 Dimensionality Reduction

Principal component analysis (PCA) (Jolliffe and Cadima, 2016) was performed on the scaled expression matrix of 3,000 HVGs. Linear dimensionality reduction by PCA is the standard first step in scRNA-seq workflows: it suppresses gene-level technical noise, decorrelates the feature space, and makes downstream computation tractable at the scale of tens of thousands of cells (Luecken and Theis, 2019; Tsuyuzaki et al., 2020). We retained 50 principal components (PCs), which is the default for this workflow and is consistent with the elbow of the scree plot (Supplementary Figure S2A), while reducing computational complexity for the models used in the analysis. Within the retained subspace, the first 20 components account for 87% and the first 30 for 93% of the captured variance (Supplementary Figure S2B); we note that these are proportions of the variance spanned by the retained components, not of total transcriptome variance. Retaining on the order of 30–50 PCs is common practice for datasets of this size, and benchmarking studies indicate that downstream results are generally robust to the precise number of components retained beyond the elbow (Luecken and Theis, 2019; Tsuyuzaki et al., 2020). Importantly, PCA preserves a linear, invertible mapping between components and genes, which is what makes the gene-level back-projection described below (Gene Importance Extraction) mathematically well defined (Jolliffe and Cadima, 2016). All random seeds were set to 42 throughout the analysis to ensure reproducibility (Sandve et al., 2013).

### 2.4 Subject-wise Train/Test Split

To prevent data leakage we strictly implemented subject-wise (grouped) splitting. Cells drawn from the same animal share technical characteristics (capture batch, sequencing lane, ambient RNA background) and biological characteristics (genotype, surgical outcome, infarct extent), and therefore violate the independence assumption underlying a random cell-wise split (Kaufman et al., 2012). Splitting repeated measurements from the same subject at the record level, rather than at the subject level, is a well-known cause of over-optimistic performance in biomedical machine learning. It can inflate apparent accuracy by a wide margin (Saeb et al., 2017; Little et al., 2017). The same problem in single-cell genomics is called pseudoreplication. It happens when individual cells, rather than animals, are treated as the unit of replication, and it strongly inflates false-discovery rates (Squair et al., 2021; Zimmerman et al., 2021). We therefore assigned every cell from a given mouse exclusively to either the training or the test set.

- **Training set:** sham1, sham2, mcao1, mcao2 (4 mice; 37,883 cells)
- **Test set:** sham3, mcao3 (2 mice; 16,716 cells)

PCA was fitted solely on training data, and test cells were then projected onto a calculated PCA transformation (using training data) called PCA space. This approach ensures complete separation between training and testing data, preventing the optimistic performance estimates that arise when cells from the same animal appear in both sets (Saeb et al., 2017; Squair et al., 2021).

### 2.5 Machine Learning Models

We evaluated eight machine learning models representing three different groups, ensuring a variety in model architecture and learning assumptions:

#### 2.5.1 Classical Machine Learning (3 models)

##### Logistic Regression (LR)

We used L2-regularized (ridge) logistic regression with regularization strength C=1.0 (Hastie et al., 2009), implemented in scikit-learn (Pedregosa et al., 2011), which provides a linear baseline model with interpretation of coefficients produced for each gene contributing to the classification.

##### Random Forest (RF)

A bootstrap-aggregated ensemble of 200 decision trees with a maximum depth of 10 (Breiman, 2001), using Gini impurity for split selection (selecting the optimal way to split the data at each node of the tree). Feature importance was calculated from the mean decrease in impurity across all trees. Because impurity-based importance is known to be biased toward continuous and high-cardinality predictors (Strobl et al., 2007), these scores were used only for relative ranking of features within a single model and were never compared in absolute terms across model families.

##### XGBoost

We used a gradient boosting model (Chen and Guestrin, 2016) with 200 trees (estimators) with a maximum depth of 6, learning rate of 0.1, and subsample ratio of 0.8. Feature importance is calculated based on how much each feature improves the model during split the data in a decision tree which is called gain-based metrics.

#### 2.5.2 Hybrid Deep Learning (1 model)

##### CNN-XGBoost

A two-stage architecture combining 1D convolutional neural networks (CNN) for feature extraction with XGBoost for classification. The CNN consists of three convolutional layers with ReLU activation and batch normalization (Ioffe and Szegedy, 2015), and learns complex (nonlinear) hierarchical feature representations from the 50-dimensional PCA embeddings (LeCun et al., 2015). These learned features are passed to XGBoost for final classification.

#### 2.5.3 Transformer-Inspired Models (4 models)

We implemented four transformer-inspired architectures (Vaswani et al., 2017), each trained from scratch on our PCA-reduced features. These models are inspired by published foundation models but trained from beginning rather than fine-tuned from pretrained weights, allowing fair comparison across different models:

##### scGPT-inspired (Cui et al., 2024)

256-dimensional embeddings, 8 attention heads, 4 transformer layers with learnable positional encodings.

##### scBERT-inspired (Yang et al., 2022)

512-dimensional embeddings, 8 attention heads, 6 transformer layers with BERT-style masked attention (Devlin et al., 2019).

##### scFormer-inspired (Cui et al., 2022)

384-dimensional embeddings, 6 attention heads, 3 transformer layers with gene-aware attention mechanisms.

##### Geneformer-inspired (Theodoris et al., 2023)

512-dimensional embeddings with rank-based input encoding, reproducing the rank-value gene expression encoding of the original Geneformer architecture.

All transformer models were trained using the AdamW optimizer (Loshchilov and Hutter, 2019) with learning rate 2 *×* 10*^−^*^4^, weight decay of 0.01, early stopping on a held-out validation split (patience = 5 epochs) to limit overfitting (Prechelt, 1998), and 10-fold stratified cross-validation on the training set (Kohavi, 1995).

### 2.6 Model Evaluation

Classification performance was evaluated on the held-out (independent) test set using the following metrics:

- Area under the receiver operating characteristic curve (AUC-ROC) (Hanley and McNeil, 1982)
- Accuracy, precision, recall, and F1 score
- Precision–recall curves, which are more informative than ROC curves when class prevalence is skewed (Saito and Rehmsmeier, 2015)
- 95% confidence intervals calculated using bootstrap resampling (1,000 iterations) (Efron and Tibshirani, 1993)

All test cells come from only two animals. The bootstrap intervals therefore measure variability within those two animals, and should not be read as confidence intervals for the wider mouse population. Generalization across subjects is limited by the number of animals, not by the number of cells (Varoquaux, 2018; Zimmerman et al., 2021).

### 2.7 Gene Importance Extraction

Gene-level importance scores were derived through model-specific approaches appropriate for each architecture:

#### Logistic Regression

Absolute values of learned coefficients, representing the magnitude of each PC’s contribution to classification.

#### Random Forest / XGBoost

Built-in feature importance metrics based on Gini impurity reduction (RF) or gain (XGBoost).

#### Transformer Models (scGPT, scBERT, scFormer)

L2 norm of the input projection layer weights, capturing how strongly each PC influences the learned representations.

For all models, PC-level importance scores were mapped to gene-level scores using the PCA loadings matrix through the following equation:

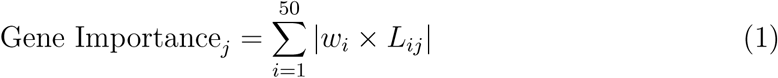

where *w_i_* is the importance score for principal component *i* and *L_ij_* is the loading of gene *j* on component *i*. This back-projection approach enables biologically interpretable gene rankings from models that operate on dimensionality-reduced representations, and is well defined precisely because the PCA mapping between components and genes is linear (Jolliffe and Cadima, 2016). These importance scores are magnitude-based and are used here for ranking and prioritisation. They identify the genes contributing most strongly to the discriminative axes, which is precisely what the consensus procedure requires. The direction of expression change is not encoded in the ranking and can be obtained from the expression data for any gene of interest. We used each model’s own importance measure rather than applying one shared explainer such as SHAP (Lundberg and Lee, 2017) to every architecture. This was deliberate. If all models were explained by the same algorithm, their agreement could come from that algorithm rather than from the models themselves (Rudin, 2019).

### 2.8 Consensus Gene Identification

To ensure robust biomarker identification that is not limited by individual model biases, we applied a multi-model consensus approach. Ensemble feature selection of this kind gives more stable and more reproducible biomarker signatures than selection from a single model (Abeel et al., 2010; Lopez-Rincon et al., 2020). The reason is that an error caused by one model’s assumptions is unlikely to be repeated by a very different model (Dietterich, 2000). From the eight trained models, six were suitable for PCA-based gene importance back-projection: Logistic Regression, Random Forest, XGBoost, scGPT-inspired, scBERT-inspired, and scFormer-inspired.

Two models were excluded from the consensus analysis for methodological reasons:

- **Geneformer-inspired:** It uses rank-based input tokenization (transforming gene expression into tokens) that fundamentally alters the relationship between input features and model weights, disrupting the direct mapping required between PCA components and genes. Hence, making PCA-based on importance back-projection mathematically inappropriate.
- **CNN-XGBoost:** The convolutional layers create nonlinear feature transformations that break the direct interpretability of PCA loadings for gene-level attribution.

Importantly, both excluded models achieved high classification performance (Gene-former AUC = 0.9778; CNN-XGBoost AUC = 0.9954), confirming their predictive validity. Their exclusion affects only the interpretability analysis, not performance evaluation.

For consensus gene identification:

1. Top 100 genes were selected from each of the 6 interpretable models
2. Genes appearing in *≥*4/6 model lists were designated as consensus genes
3. Consensus genes were ranked by total model vote count

Both the list depth (top 100) and the agreement threshold (*≥*4/6) are free parameters. Any such threshold is a trade-off between selection stability and sensitivity (Nogueira et al., 2018). We therefore report the full distribution of agreement levels across all six models (Supplementary Figure S4), so the effect of the threshold can be seen directly.

### 2.9 Functional Enrichment Analysis

Gene Ontology (GO) (Ashburner et al., 2000) and KEGG (Kanehisa and Goto, 2000) pathway enrichment analyses were performed using clusterProfiler v4.18 (Wu et al., 2021) with the following parameters:

- Background gene set: All 3,000 highly variable genes
- Statistical test: Hypergeometric test
- P-value threshold: 0.05
- Multiple testing correction: Benjamini–Hochberg false discovery rate (FDR) (Benjamini and Hochberg, 1995)
- Minimum gene set size: 10; Maximum gene set size: 500

Identical parameters and the identical background were used for the GO and KEGG analyses, so that the two are directly comparable.

We restricted the background to the 3,000 highly variable genes rather than the full transcriptome. This is because the consensus genes could only have been selected from that set. Using an unrestricted background in over-representation analysis is a known cause of inflated enrichment significance (Huang et al., 2009; Timmons et al., 2015).

## 3 Results

### 3.1 Study Overview

We developed a multi-model consensus framework to identify robust stroke biomarkers from single-cell transcriptomic data (Figure 1). Starting with scRNA-seq data from a mouse MCAO model, we applied rigorous quality control, implemented subject-wise train/test splitting, and trained eight machine learning models covering classical, hybrid, and transformer-inspired architectures. Consensus genes were identified based on agreement across six interpretable models.

**Figure 1:**
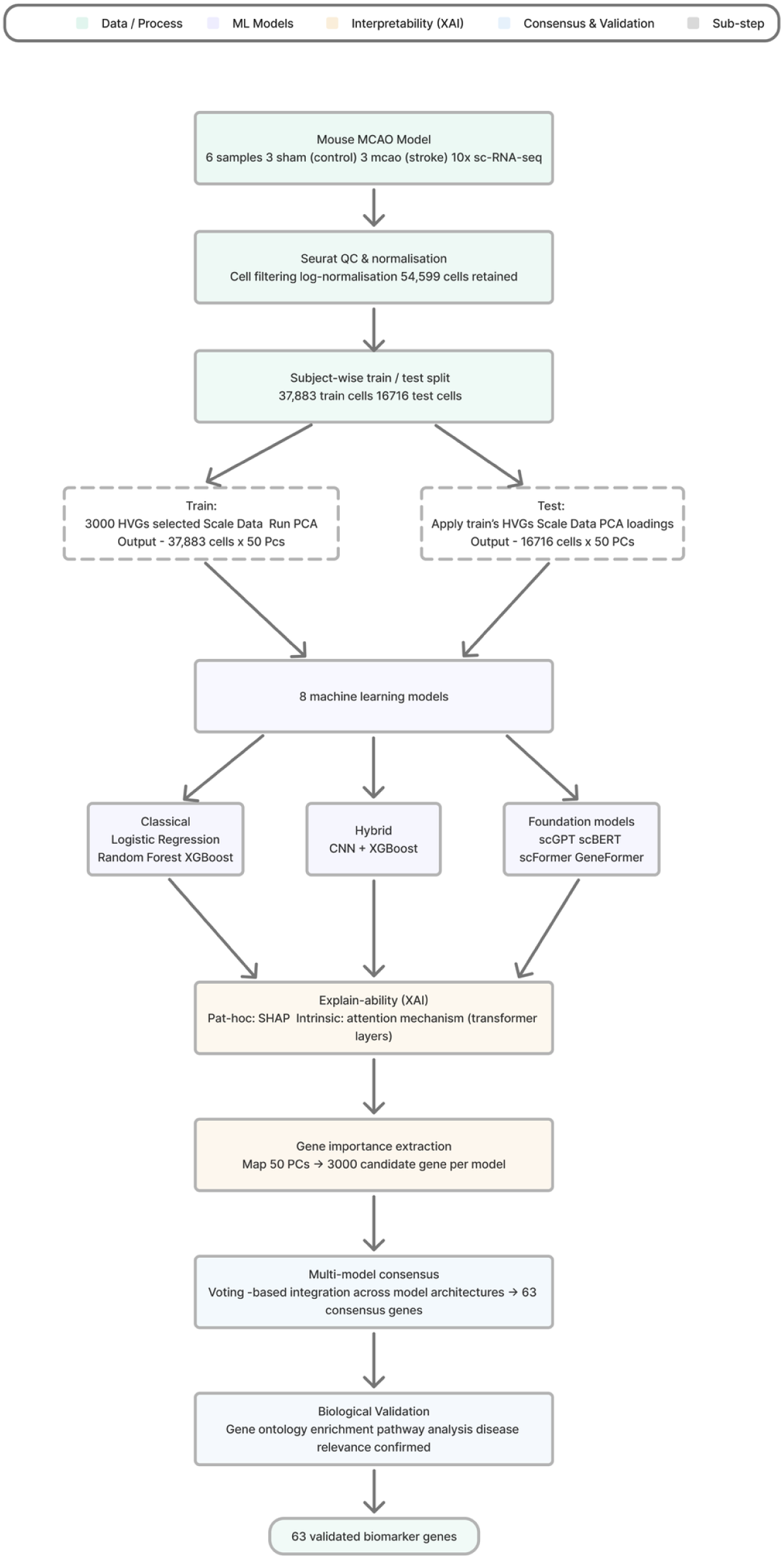
Multi-model consensus framework for stroke biomarker discovery. Overview of the analysis pipeline from single-cell RNA sequencing through quality control, PCA-based feature extraction, eight-model training with subject-wise splitting, gene importance back-projection, and consensus identification. The framework integrates classical ML, hybrid deep learning, and transformer-inspired models.

### 3.2 Data Characteristics and Quality Control

After stringent quality control filtering, our dataset comprised 54,599 high-quality single cells (Table 1). The training set contained 37,883 cells from four mice (two sham, two MCAO), while the independent (held-out) test set contained 16,716 cells from two mice (one sham, one MCAO). This subject-wise splitting ensures that model performance reflects generalization to completely unseen biological samples rather than learning patterns within known samples (Saeb et al., 2017; Zimmerman et al., 2021).

**Table 1:** Sample characteristics after quality control filtering. Subject-wise splitting ensures no data leakage between training and test sets.

| Sample | Condition | Set | Cells |
| --- | --- | --- | --- |
| sham1 | Control | Training | 8,410 |
| sham2 | Control | Training | 8,144 |
| mcao1 | MCAO | Training | 10,801 |
| mcao2 | MCAO | Training | 10,528 |
| **Training Total** |  |  | **37,883** |
| sham3 | Control | Test | 9,508 |
| mcao3 | MCAO | Test | 7,208 |
| **Test Total** |  |  | **16,716** |
| **Grand Total** |  |  | **54,599** |

### 3.3 Model Performance

All eight machine learning models achieved excellent classification performance on the held-out test set, with AUC values ranging from 0.9778 to 0.9975 (Table 2, Figure 2). The scGPT-inspired transformer achieved the highest AUC (0.9975, 95% CI: 0.9970–0.9980), followed closely by logistic regression (AUC = 0.9968, 95% CI: 0.9963–0.9972). Notably, the simplest model (logistic regression) performed comparably to the most complex architectures, suggesting that the biological signal distinguishing stroke from control cells is sufficiently strong to be captured by linear relationships. This is consistent with recent systematic evaluations reporting that deep architectures do not yet reliably outperform simple linear baselines on single-cell prediction tasks (Ahlmann-Eltze et al., 2025).

**Figure 2:**
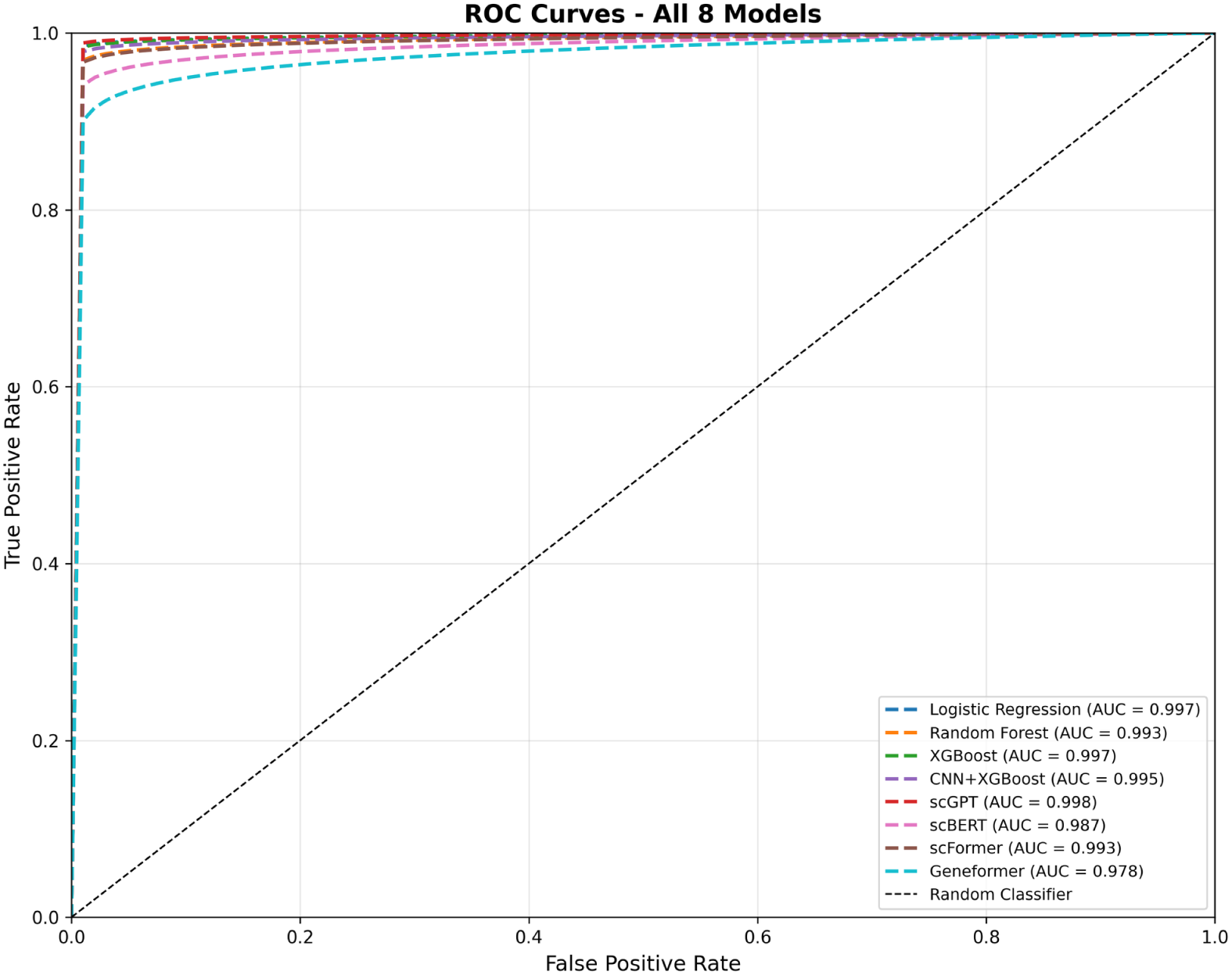
Receiver operating characteristic curves for all eight models. All models achieved AUC *>*0.97 on the held-out test set (n=16,716 cells from 2 mice). Classical ML models shown in blue/orange/green; hybrid CNN+XGBoost in purple; transformer-inspired models (scGPT, scBERT, scFormer, Geneformer) in red/pink/brown/cyan. Diagonal dashed line indicates random classifier performance.

**Table 2:** Classification performance of all eight machine learning models on the held-out test set. Models are ranked by AUC. All models included in consensus analysis unless otherwise noted.

| Model | AUC | 95% CI | F1 | Consensus |
| --- | --- | --- | --- | --- |
| scGPT-inspired | 0.9975 | [0.9970–0.9980] | 0.953 | Yes |
| Logistic Regression | 0.9968 | [0.9962–0.9974] | 0.949 | Yes |
| XGBoost | 0.9966 | [0.9961–0.9971] | 0.932 | Yes |
| CNN-XGBoost | 0.9954 | [0.9946–0.9961] | 0.927 | No<sup>a</sup> |
| Random Forest | 0.9932 | [0.9924–0.9941] | 0.918 | Yes |
| scFormer-inspired | 0.9927 | [0.9915–0.9939] | 0.946 | Yes |
| scBERT-inspired | 0.9869 | [0.9853–0.9886] | 0.934 | Yes |
| Geneformer-inspired | 0.9778 | [0.9756–0.9798] | 0.919 | No<sup>b</sup> |

<sup>a</sup>CNN feature extraction incompatible with PCA back-projection
<sup>b</sup>Rank-based tokenization incompatible with PCA back-projection

The high performance across all model families—from the simplest linear classifier to complex transformer architectures—suggests that stroke-associated transcriptional changes create robust separability (stroke cells from control cells) in the data. This consistency is itself an informative result. It shows that the transcriptional response to ischemia at 24 hours is strong enough to be recovered by models built on very different assumptions. Because all eight models fall within a narrow performance range, we interpret the comparison as evidence of a reproducible biological signal rather than as a ranking of architectures. The 10-fold cross-validation results (Supplementary Figure S3) confirmed stable performance across training folds with minimal variance.

### 3.4 Gene Signature Convergence Across Model Families

Analysis of gene importance rankings revealed distinct clusters of agreement among model families (Figure 3). Classical ML models (logistic regression, random forest, XGBoost) showed moderate pairwise overlap, quantified as the Jaccard index between top-100 gene lists—a standard measure of feature-selection stability (Nogueira et al., 2018)—of 0.21–0.83, while transformer-inspired models demonstrated remarkably high convergence (0.98–1.00), suggesting they capture similar biological features despite architectural differences.

**Figure 3:**
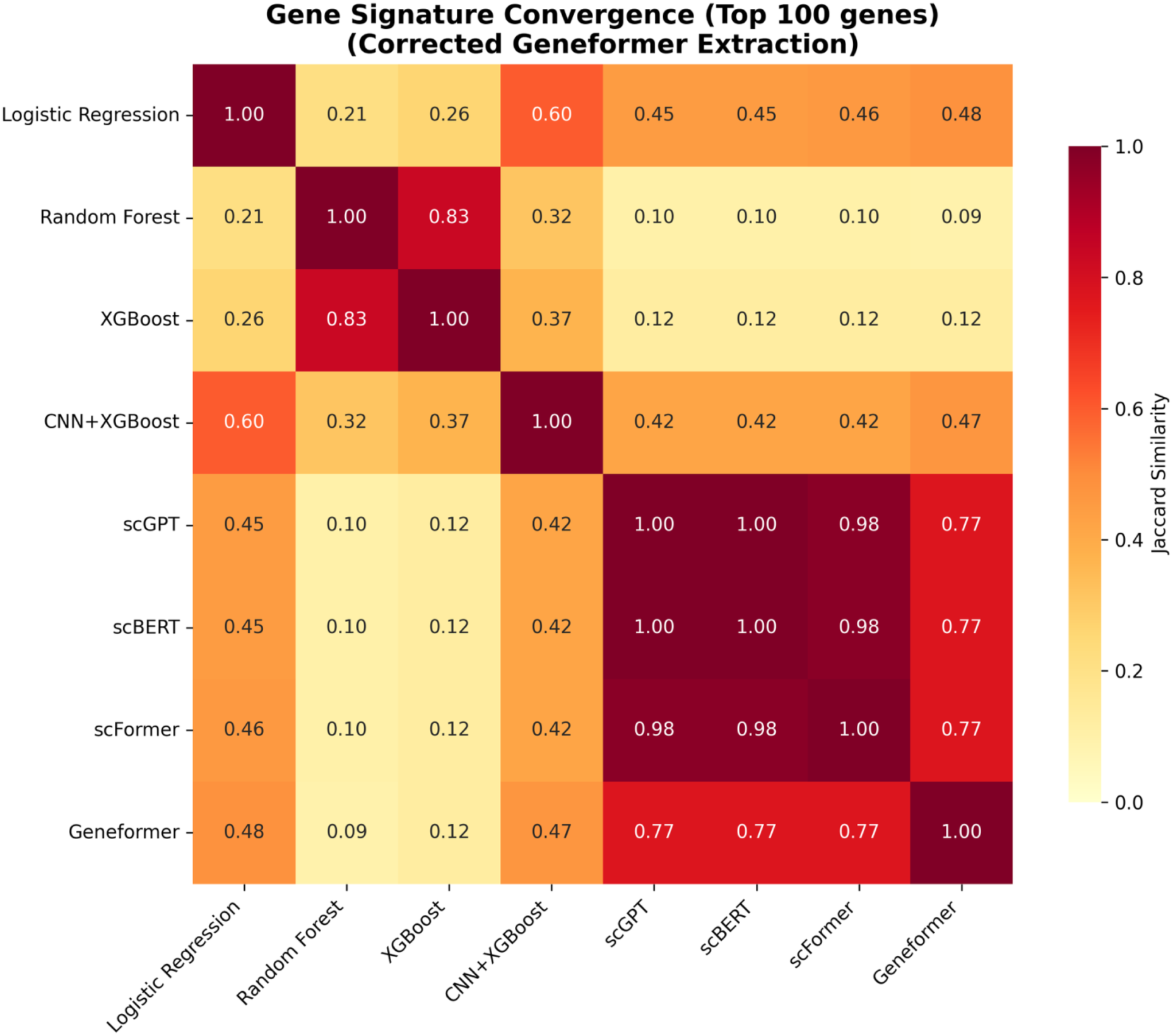
Gene signature convergence across all eight models. Heatmap showing pairwise Jaccard similarity of top-100 gene lists between models. Classical ML models (Logistic Regression, Random Forest, XGBoost) form one cluster with moderate internal similarity (21–83%). Transformer-inspired models (scGPT, scBERT, scFormer, Gene-former) form a highly convergent cluster (77–100%). CNN+XGBoost occupies a unique position with 60% overlap with Logistic Regression and 42–47% overlap with foundation models. The high inter-correlation among transformers justified limiting their representation in the consensus to avoid over-weighting one model family.

The high inter-correlation among transformer models (Figure 3, lower-right quadrant) justified our decision to limit transformer representation in the consensus analysis. Including all four transformer models would have given too much weight to the transformer model family in the voting process. Additional analysis confirmed that Gene-former showed 77% overlap with scGPT, scBERT, and scFormer, further supporting this decision.

### 3.5 Consensus Gene Identification

Multi-model consensus analysis identified 63 genes appearing in the top-100 lists of at least four of six interpretable models (threshold: *≥*4/6; Supplementary Table S1). Seventeen genes achieved perfect 6/6 agreement across all models, representing the highest-confidence biomarker candidates (Table 3). Two of these unanimously selected genes, Spp1 and Lpl, are core markers of the disease-associated microglia (DAM) program reported in neurodegeneration and acute brain injury (Keren-Shaul et al., 2017; Hammond et al., 2019). This gives independent literature support for the consensus signature, and none of it was used when deriving the signature.

**Table 3:** Top consensus genes with unanimous or near-unanimous model agreement. Function annotations based on Gene Ontology and literature review.

| Rank | Gene | Models | Function |
| --- | --- | --- | --- |
| 1 | Spp1 | 6/6 | Osteopontin; macrophage activation, tissue remodeling |
| 2 | Cdkn1a | 6/6 | p21; cell cycle arrest, senescence |
| 3 | Cd24a | 6/6 | Cell surface glycoprotein; immune modulation |
| 4 | Il1rn | 6/6 | IL-1 receptor antagonist; anti-inflammatory (Anakinra target) |
| 5 | Ifitm6 | 6/6 | Interferon-induced transmembrane protein; antiviral response |
| 6 | H2-Eb1 | 6/6 | MHC Class II; antigen presentation |
| 7 | H2-Aa | 6/6 | MHC Class II; antigen presentation |
| 8 | Fth1 | 6/6 | Ferritin heavy chain; iron storage, oxidative stress |
| 9 | Lpl | 6/6 | Lipoprotein lipase; lipid metabolism |
| 10 | Anxa1 | 6/6 | Annexin A1; anti-inflammatory, resolution of inflammation |
| 11 | Adam8 | 6/6 | Metalloprotease; cell adhesion, inflammation |
| 12 | Ch25h | 6/6 | Cholesterol 25-hydroxylase; innate immunity |
| 13 | Plac8 | 6/6 | Placenta-specific 8; immune cell marker |
| 14 | Napsa | 6/6 | Napsin A; antigen processing |
| 15 | Cd72 | 6/6 | B-cell surface marker; immune regulation |
| 16 | Tubb6 | 6/6 | Beta-tubulin 6; cytoskeleton |
| 17 | Sirpb1c | 6/6 | <sup>18</sup>Signal-regulatory protein; immune modulation |

Several of the consensus genes have direct therapeutic relevance. Il1rn (6/6 models) encodes the IL-1 receptor antagonist, a natural inhibitor of the pro-inflammatory IL-1*β* pathway. Its recombinant form, Anakinra, is FDA-approved for rheumatoid arthritis and has completed Phase II trials in ischemic stroke with a good safety profile and a reduction in systemic inflammatory markers, though without a significant effect on functional outcome (Smith et al., 2018; Emsley et al., 2005; Sobowale et al., 2016). Plat (5/6 models) encodes tissue plasminogen activator, the molecular target of Alteplase, the primary FDA-approved thrombolytic therapy for acute stroke. Spp1 and Fth1 (both 6/6) act on tissue repair and cell death. Osteopontin (Spp1) is strongly induced around the infarct, where it takes part in glial scar formation and debris clearance by macrophages (Ellison et al., 1998), while ferritin heavy chain (Fth1) stores iron, and its degradation releases iron into the labile pool that drives ferroptotic cell death after ischemia (Dixon et al., 2012; Shi et al., 2025). Anxa1 (6/6) is a key mediator of inflammation resolution, and its synthetic peptide Ac2-26 is neuroprotective in preclinical stroke models (Li et al., 2021; Perretti and D’Acquisto, 2009). None of these genes was chosen in advance; all emerged from the consensus.

### 3.6 A Novel Dual-Interferon Gene Signature

Twelve of the 63 consensus genes are annotated as interferon-responsive, and the way they distribute across the Type I and Type II response terms defines what we describe as the “dual-interferon signature” (Figure 4). Response to interferon-beta (GO:0035456) contains 10 of the consensus genes and response to type II interferon (GO:0034341) contains 7. Critically, five genes are annotated to *both* terms, so the two arms of the signature overlap rather than being disjoint. Gene groups below are assigned strictly by Gene Ontology membership; model vote counts are given in parentheses.

**Figure 4:**
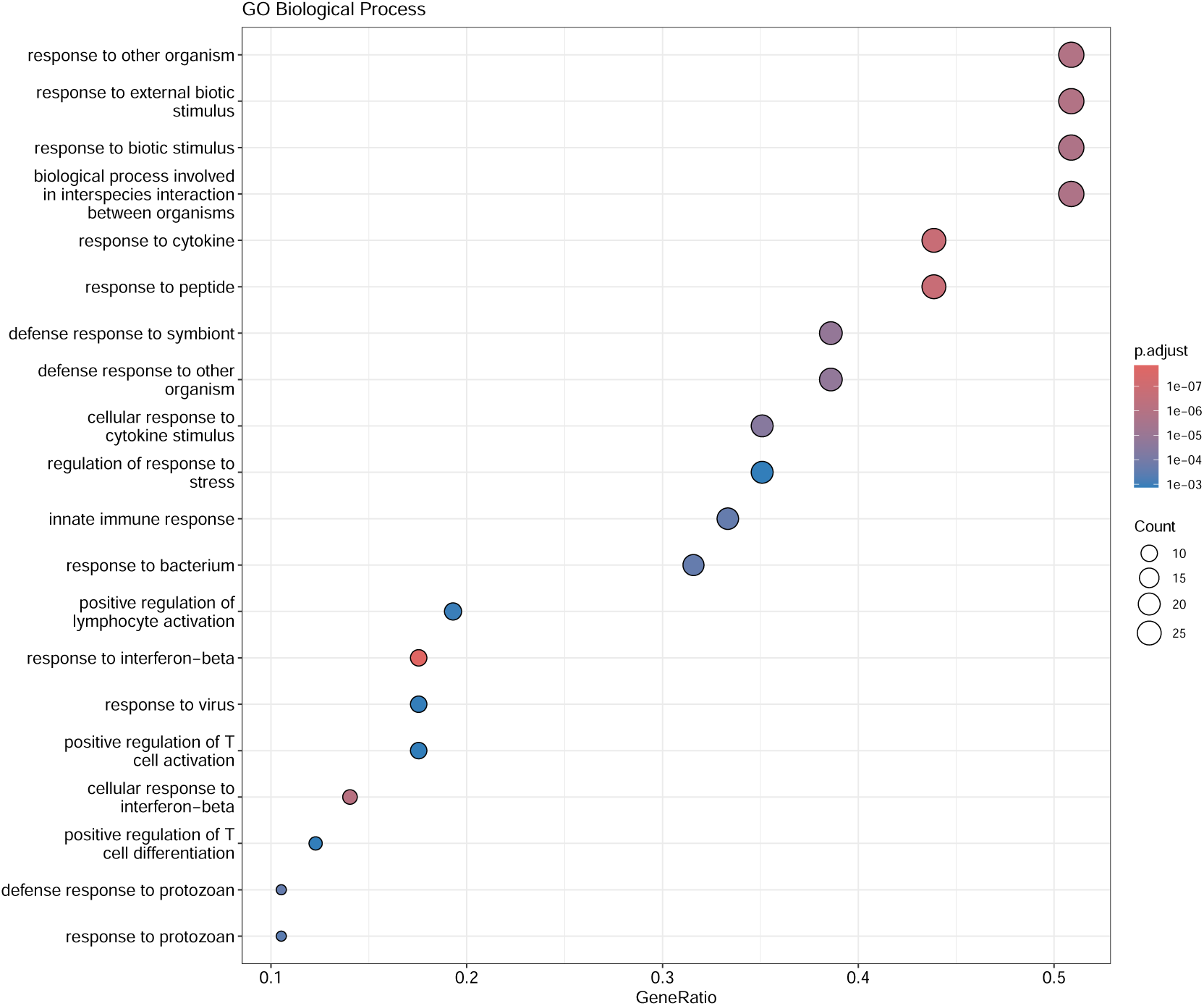
Gene Ontology enrichment analysis of 63 consensus genes. Dot plot showing the most significantly enriched biological processes, tested against a background of the 3,000 highly variable genes from which the consensus was drawn. Point size indicates the gene ratio (proportion of consensus genes in each term); color represents adjusted p-value (FDR). Type I and Type II interferon response terms dominate the profile, with MHC class II antigen presentation also significant.

**Responsive to both Type I and Type II interferon (5 genes):**

- Ifitm6 (6/6), Ifitm1 (4/6) — interferon-induced transmembrane proteins (Bailey et al., 2014)
- Gbp2 (4/6) — guanylate-binding protein (Kim et al., 2012)
- Igtp (4/6), Tgtp1 (4/6) — immunity-related GTPases (MacMicking, 2012)

**Type I interferon (IFN-*β*) responsive only (5 genes):**

- Ifit3 (4/6) — interferon-induced protein with tetratricopeptide repeats
- Ifi47 (4/6), Ifi205 (4/6) — interferon-inducible proteins
- Iigp1 (4/6), Tgtp2 (4/6) — immunity-related GTPases (MacMicking, 2012)

**Type II interferon (IFN-*γ*) responsive only (2 genes):**

- Gbp4 (4/6) — guanylate-binding protein (Kim et al., 2012)
- Cd74 (4/6) — MHC class II invariant chain

The five dual-responsive genes form the core of the signature. These genes are induced by both the IFNAR and IFNGR receptor systems. Because at least four of six independent models selected each of them, both interferon pathways appear to be active at the same time after stroke, rather than one being inferred from the other. Ifitm6 was selected by all six interpretable models.

These findings fit with evidence that both interferon types play important and sometimes opposing roles after stroke (Deczkowska et al., 2016). Type I interferons (IFN-*α*/*β*) are induced rapidly after ischemic injury and can be either protective or harmful depending on timing and cell type (Iadecola and Anrather, 2011). Type II interferon (IFN-*γ*) comes mainly from infiltrating T cells and NK cells; it pushes microglia toward a pro-inflammatory state and worsens infarct size in experimental stroke (Yilmaz et al., 2006; Iadecola et al., 2020). The dual signature likely reflects the recruitment of different interferon-producing cells over time, plasmacytoid dendritic cells for Type I and T cells for Type II, in line with the known sequence of immune cell entry into the ischemic hemisphere (Gelderblom et al., 2009; Liesz et al., 2009), followed by activation of shared interferon-stimulated genes downstream of both receptors (Deczkowska et al., 2016; MacMicking, 2012).

### 3.7 Pathway Enrichment Analysis

Gene Ontology enrichment analysis of the 63 consensus genes, against a background restricted to the 3,000 highly variable genes, identified 101 significantly enriched biological processes, 12 molecular functions and 16 cellular components (FDR *<* 0.05; Figure 4). Of the 63 genes, 57 mapped to Entrez identifiers and were used for testing. The enrichment profile is dominated by interferon response:

- Response to interferon-beta, i.e. Type I interferon (GO:0035456, FDR = 1.4*×*10*^−^*^8^; 10/57 genes)
- Cellular response to interferon-beta (GO:0035458, FDR = 7.3*×*10*^−^*^7^; 8/57 genes)
- Response to type II interferon, i.e. IFN-*γ* (GO:0034341, FDR = 4.5*×*10*^−^*^3^; 7/57 genes)
- Antigen processing and presentation via MHC class II (GO:0019886, FDR = 1.0*×*10*^−^*^2^; 4/57 genes)
- Inflammatory response (GO:0006954, FDR = 1.5*×*10*^−^*^2^; 16/57 genes)
- Defense response to virus (GO:0051607, FDR = 2.0*×*10*^−^*^2^; 7/57 genes)

Notably, response to interferon-beta is the single most significantly enriched process in the entire analysis, and both Type I and Type II interferon terms remain significant against this conservative background. The dual-interferon signature is therefore not an artefact of a permissive gene universe.

KEGG over-representation analysis was performed on the same gene set against the same background. No KEGG pathway survived Benjamini–Hochberg correction (122 pathways tested; smallest adjusted p-value 0.162). Twenty-one pathways reached nominal significance (*p <* 0.05), but ten of these were driven entirely by the same three MHC class II genes (H2-Aa, H2-Ab1, H2-Eb1), which are annotated to a large number of unrelated KEGG disease maps. We therefore report Gene Ontology as the primary enrichment analysis and treat the KEGG result as negative. Complete per-pathway statistics are provided in the accompanying code repository.

## 4 Discussion

We present a multi-model machine learning consensus framework that identifies 63 robust stroke biomarker candidates from a methodologically diverse set of eight algorithms. All models achieved high classification performance (AUC 0.9778–0.9975), and a dual-interferon gene signature appeared consistently across model families. Therapeutically actionable targets with existing FDA-approved drugs, notably Il1rn (Anakinra) and Plat (tPA), were identified without any prior assumption about stroke biology. Interferon signaling and IL-1 pathway activation are well described in ischemic stroke (Iadecola et al., 2020; Sobowale et al., 2016), and their independent recovery here is strong evidence that the framework finds genuine biology. Building on that foundation, the analysis identifies a pattern in which five consensus genes respond to both Type I and Type II interferon.

The central insight is that genes chosen by a single model may reflect that model’s own patterns rather than true biology. The close agreement between simple linear models and complex transformers shows that the main transcriptional difference between stroke and control cells can be detected by very different mathematical frameworks. This convergence gives confidence that the 63 consensus genes represent real biology, consistent with prior work showing that ensemble feature selection improves the reproducibility of biomarker signatures (Abeel et al., 2010; Dietterich, 2000). The high inter-correlation among the transformer models (Jaccard *>*0.98) means they give overlapping rather than independent evidence (Dietterich, 2000; Nogueira et al., 2018), and we limited their representation in the consensus to three of six models for this reason.

Three design choices shaped the analysis. The transformer models were trained from scratch on PCA-reduced features rather than fine-tuned from pretrained weights. This holds the input representation constant across all eight architectures, so that differences in gene selection reflect the models rather than their pretraining data; we use the “-inspired” suffix throughout to make this explicit. PCA was chosen because its linear loading structure allows model importance to be back-projected exactly onto genes (Jolliffe and Cadima, 2016; Tsuyuzaki et al., 2020), an operation that nonlinear embeddings such as variational autoencoders (Lopez et al., 2018) do not support. The subject-wise split prevents data leakage and gives a true test of generalization to unseen animals. The cost is a test set of only two mice, so the relevant uncertainty is between subjects rather than between cells (Varoquaux, 2018).

Several directions would extend this work. Validation in human stroke cohorts is the priority, given the high failure rate of preclinical-to-clinical translation in stroke and the STAIR call for replication across species and models (O’Collins et al., 2006; Fisher et al., 2009). Larger animal numbers would give more reliable estimates between animals, since power depends on the six mice rather than the 54,599 cells (Squair et al., 2021; Zimmerman et al., 2021). Dedicated doublet detection (McGinnis et al., 2019; Wolock et al., 2019), ambient-RNA correction (Young and Behjati, 2020) and batch-effect correction (Tran et al., 2020) could be added to the pipeline. Enrichment results are sensitive to the background set and to annotation bias (Huang et al., 2009; Timmons et al., 2015) and should be read as hypothesis-generating until confirmed experimentally. Extending the analysis to multiple timepoints and to individual cell types would show how the signature evolves across the phases of stroke. Pretrained foundation models with gene-level tokenization are a natural next step for the framework, and the relevance of Il1rn suggests an immediate opportunity to re-analyze existing Anakinra trial data stratified by IL1RN expression.

## 5 Conclusions

Multi-model machine learning consensus provides a robust approach to biomarker discovery that reduces dependence on any single algorithm. Its strength lies in the convergence of methodologically distinct models on the same gene set, which provides a level of confidence that no individual model can supply on its own. Applied to ischemic stroke single-cell transcriptomics, this framework identified 63 consensus genes with strong biological relevance, including a novel dual-interferon signature and therapeutically actionable IL-1 pathway components. The agreement of classical, ensemble, and deep learning methods on these genes supports their validity as stroke biomarkers and provides a prioritized foundation for translational research.

## Supporting information

https://drive.google.com/file/d/1UTPgZEj587PtPuuhl9b51ljJQXIdZenc/view?usp=drivesdk

## Data Availability Statement

The original single-cell RNA sequencing data are publicly available in the Gene Expression Omnibus under accession number **GSE174574**. All analysis code and processed data supporting this study are available at https://github.com/AliceCyrus/ multimodel-consensus-stroke-scrnaseq.

## Author Contributions

AS and AC conceived and designed the study. AS performed all computational analyses including data preprocessing, model development, and statistical validation. AC provided biological interpretation and clinical context. Both authors contributed to manuscript writing and approved the final version.

## Acknowledgments

The authors thank Md Kashif Akram for technical guidance and valuable discussions during the computational analysis phase.

