## Supplementary material for "Multi-Model Machine Learning Consensus Identifies a Dual-Interferon Gene Signature in Ischemic Stroke: An Integrative Single-Cell Transcriptomic Analysis": https://drive.google.com/file/d/1UTPgZEj587PtPuuhl9b51ljJQXIdZenc/view?usp=drivesdk

### Supplementary Figures

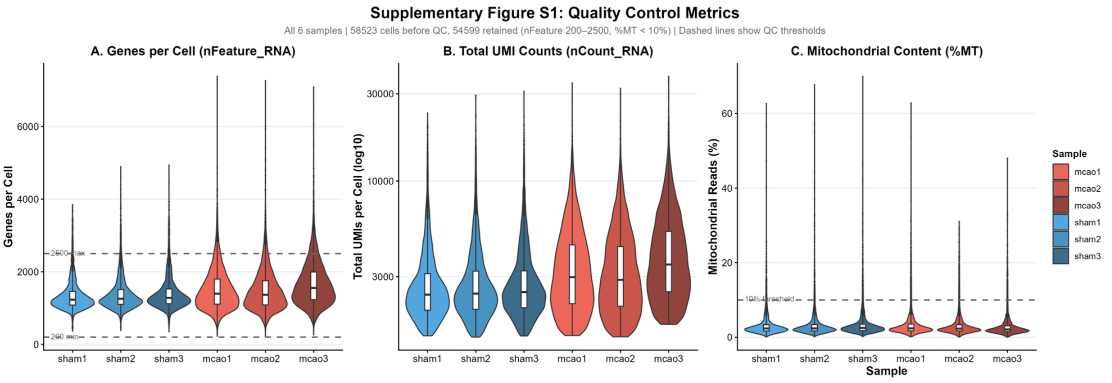


Figure S1: **Quality control metrics.** Distribution of (A) genes detected per cell, (B) total UMIs per cell, and (C) mitochondrial gene percentage for all 58,523 cells before filtering, shown per sample. Dashed lines mark the QC thresholds. The thin upper tails above the 2,500-gene and 10% lines are the cells removed; 54,599 cells were retained.

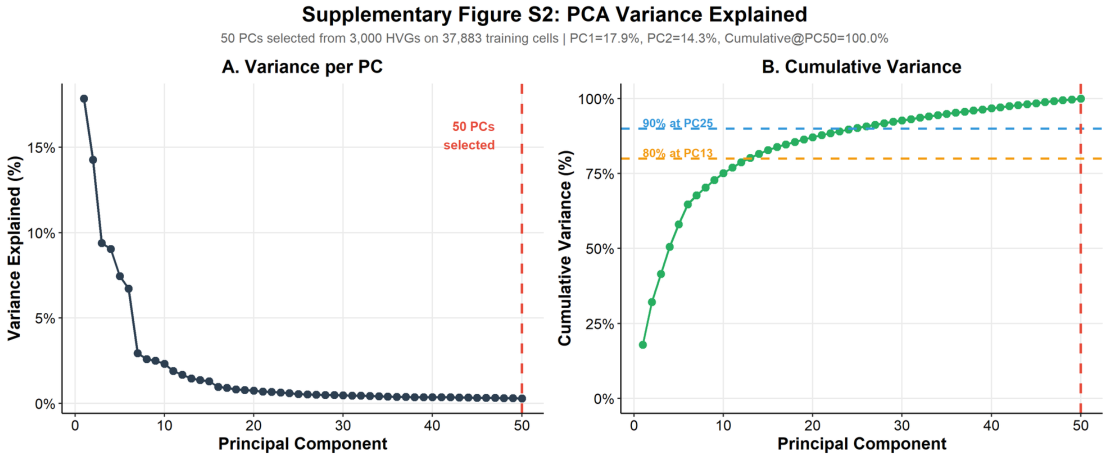


Figure S2: **PCA variance analysis.** (A) Elbow plot showing variance explained by each principal component. (B) Cumulative variance across the 50 retained components, normalized to the variance those components span, so the curve reaches 100% at PC 50; the first 30 components account for 93% of it.

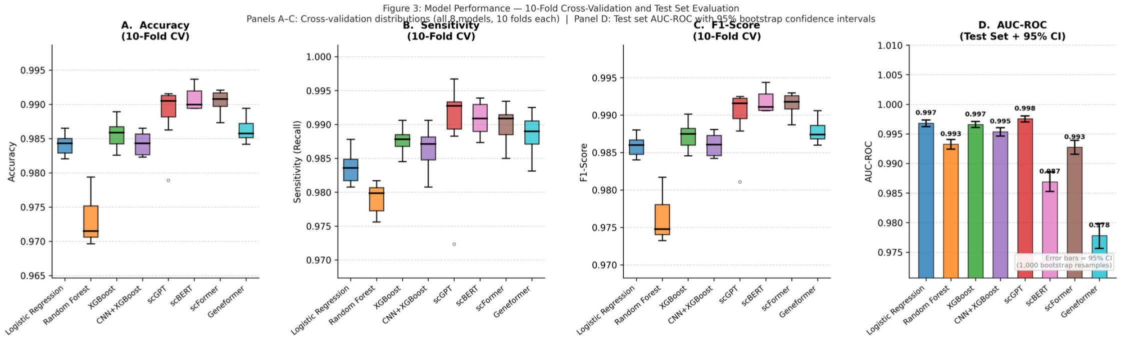


Figure S3: **Cross-validation performance stability.** Distribution of performance metrics across 10-fold stratified cross-validation for each model, demonstrating consistent performance with minimal inter-fold variance.

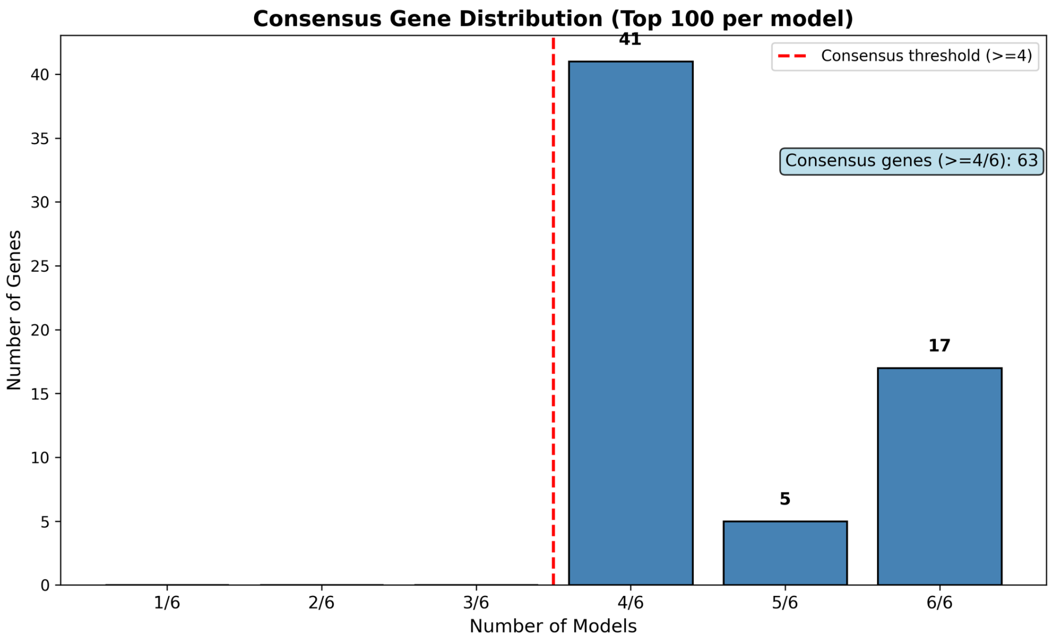


Figure S4: **Consensus gene distribution.** Number of genes achieving each level of model agreement across all six interpretable models. Bars at 1–3 models (red region, left of dashed line) contain zero genes; all consensus candidates achieved ≥4/6 agreement. The threshold of ≥4/6 identifies 63 consensus genes (41 at 4/6, 5 at 5/6, 17 at 6/6), balancing sensitivity and robustness.
